# Identification of sialic acid linkage profiles in primary liver cancers

**DOI:** 10.1101/2025.11.20.689542

**Authors:** Shaaron Ochoa-Rios, James W. Dressman, Peggi M. Angel, Richard R. Drake, Anand S. Mehta

**Affiliations:** Department of Cell and Molecular Pharmacology, Medical University of South Carolina, Charleston, South Carolina

**Author notes:** **Corresponding Author:** Anand S. Mehta, Medical University of South Carolina, 173 Ashley Avenue, Charleston, SC 29425.

**Keywords:** N-glycans, sialic acid, linkages, MALDI-IMS

## Abstract

Cholangiocarcinoma (CCA), also known as bile duct cancer, is the second most common form of primary liver cancer after hepatocellular carcinoma (HCC). Subtypes of CCA are diagnosed according to the anatomic location within the biliary tree, such as intrahepatic (iCCA) or extrahepatic (eCCA). Early diagnosis remains difficult due to the low sensitivity and specificity of current biomarkers. There has been significant interest in finding new biomarkers of CCA and much work has been performed to both discover and validate these new markers. In our previous work, we identified specific alterations in N-linked glycosylation that were associated with iCCA, which differed from non-malignant cholestasis disease and HCC. However, that analysis focused only on N-glycan structures devoid of sialic acid, as these N-glycan residues are not stable and were intentionally removed to minimize variability. Here, we explore the specificity of sialic acid residues in liver cancers, since sialic acid residues have been heavily linked to roles in tumor progression and metastasis by helping evade immunological surveillance. To circumvent the instability of sialic acid, we utilize a novel sialic acid stabilization method coupled with Matrix-Assisted Laser Desorption Ionization (MALDI) Imaging Mass Spectrometry (IMS) for detailed analysis of sialic acid linkages in iCCA tissue. We report a clear association between the linkages type of sialic acid residues with a HCC or CCA, including differentiation between CCA subtypes. Overall, we stabilize N-glycans with sialic acid residues to determine their linkages and elucidate their importance in different types of primary liver cancers and their subtypes by MALDI-IMS.

## Introduction

Cholangiocarcinoma (CCA) is a highly lethal and rare epithelial cancer arising in the biliary ducts and is the second most common type of liver cancer after hepatocellular carcinoma (HCC) [1-7]. CCA frequently consists of small nests of epithelial cancer cells surrounded by dense stromal regions of cancer-associated fibroblasts, immune cell populations, and extracellular matrix. In addition, glandular formations are also a histological characteristic of CCA [8]. CCA is classified based on the anatomic location into intrahepatic CCA (iCCA) or extrahepatic CCA (eCCA). eCCA is further classified as perihilar CCA (pCCA), or distal (dCCA) subtypes [8]. In the USA, 10–20% of cases are iCCA, 50–60% are pCCA, and 20–30% are dCCA. Increases in mortality in recent years have largely been driven by iCCA, with mortality rates from the other forms either remaining constant or falling over 20 years [9]. Indeed, the mortality rate of iCCA in the USA in 2012 was 8.8-fold higher with iCCA than with eCCA [9]. Risk factors for both iCCA and eCCA include underlying bile duct disease in the form of Carroli disease, choledochal cysts, and primary sclerosing cholangitis (PSC) [10,11]. Other factors include fluke infection, obesity, smoking, chronic viral hepatitis, and cirrhosis [3,9,12]. Of all the risk factors, the most common risk factor in Western countries is PSC while liver fluke infection is associated with CCA in Southeast Asia [13].

Currently, carbohydrate antigen 19-9 (CA19-9) is the main biomarker used for the diagnosis of CCA; however, the use of this glycoprotein has faced challenges of low specificity and sensitivity often falsely elevated in the setting of cholestasis without malignancy. A definitive diagnosis of CCA at an early stage continues to be challenging. Some of the main challenges with CCA diagnosis are that it is historically misdiagnosed as HCC (specifically for iCCA) [3] and the need for a combination of diagnostic methods including liver biopsy, and several ERCPs (endoscopic retrograde cholangiopancreatography) to establish a definitive diagnosis. Overall, there is an urgent need for the identification of diagnostic and prognostic biomarkers for CCA and its subtypes [10].

Glycosylation is one of the most common port translational modifications and is based on the addition of glycan structures on macromolecules, mainly proteins and lipids. Glycosylation has many biological roles including cell-cell communication, assisting in protein folding, and receptor signaling. At the same time, glycosylation has been linked to have many roles in cancer-related processes. Specifically, alterations in N-linked glycosylation have been reported by us and others to be strongly associated with many different types of cancer. Sialylation (addition of a sialic acid residue) and fucosylation (addition of a fucose residue) have been reported as major modifications to the N-glycan structure with the ability to correlate and distinguish between disease states. Sialylation is catalyzed by sialyltransferases (STs), each with different linkage specificity on the N-glycan, this study focuses on the terminal decoration of sialic acid residues linked at the α2-3 and/or α2-6 [14,15]. N-glycan sialyation is a modification that plays an important role in the immune system and is known to occur in cancer, indeed, CA19-9 is a sialylated glycan. Sialic acid linkages, specifically α2-3 and α2-6, have been reported to play distinct roles in various disease processes, including cancer [16-19]. However, the characterization of sialic acid residues in N-glycan structures using Matrix Assisted Laser Desorption (MALDI) as an ionization source has presented limitations due to the lability of sialic acid residues during ionization. In addition, the identification of the type of sialic acid linkages on N-glycans (α2,3 and α2,6) is also limited by mass spectrometry since the different linkages do not result in mass differences [20-22]. A sialic acid stabilization and derivatization method (alkyne-amidation eXpanded linker (AAXL)) developed by our group addresses these limitations and targets α2,3- and α2,6-linked sialic acids isomers with biorthogonal chemical labeling probes and allows for the identification of these isomers by mass spectroscopy. This method is coupled to previously established N-glycan mass spectrometry imaging protocols and allows for the detection and identification of sialic acid isomers on tissue [22-25].

## Materials and methods

### Tissue Microarrays

Tissue microarray 1 (TMA; #LV2081, Biomax, Inc) contained 208 cores with 103 cases (duplicated cores per case): consisting of fifty HCC, twenty iCCA, one clear-cell carcinoma cyst, five metastatic HCC (spleen, chest wall, cerebrum, costal bone, and lymph node), two hepatic cyst, eight tissues with cirrhosis and dysplastic nodules, ten hepatitis-infected tissues, two adjacent normal tissues (labeled here as other), and six independent normal tissues. Each core was 1.0mm in diameter and diagnosed by pathology. Tissue microarray 2 (TMA; #GA802a, Biomax, Inc) contained 80 cores with 80 cases: consisting of forty-eight cases of extrahepatic biliary duct adenocarcinoma CCA (eCCA), twenty-seven iCCA, and five adjacent normal liver tissues. Each core was 1.5mm in diameter and diagnosed by pathology. H&E staining of TMA 1 and 2 are provided in Supplementary Figures 1a and 1b, respectively.

### Enzymes and reagents

Recombinant peptide N-glycosidase F (PNGase F) PRIME (N-Zyme Scientifics), α-cyano-4-hydroxycinnamic acid (CHCA), Trifluoroacetic acid, and Harris-modified hematoxylin (Sigma-Aldrich).

### Tissue Preparation for MALDI-IMS

Unstained FFPE TMA slides were processed using established imaging workflows of MALDI-IMS for N-glycans as previously described[24]. In short, tissues were heated at 60C for 1 house, deparaffinized (with xylene washes) and rehydrated (with water and ethanol washes). TMA1 was processed for sialic acid stabilization and derivatization following a previously established method, AAXL (alkyne-amidation eXpanded linker)[22]. The AAXL method is based on the amidation-amidation reaction with dimethylamine and propargylamine. TMA 2 was processed for antigen retrieval and placed in a decloaker for 30 minutes using citraconic anhydride (Thermo Fisher Scientific) as the buffer. Both TMAs were processed for N-glycan release, PNGase F PRIME (N-[26]Preparation System (HTX Technologies, LLC). After the spray of the enzyme, slides were incubated in a humidified chamber for 2 hours at 37°C. Finally, slides are sprayed with matrix using CHCA (0.042 g CHCA in 6 mL 50% acetonitrile/49.9% water/0.1% trifluoroacetic acid) and using the same M5 TM-Sprayer as before.

### N-glycan Imaging Using MALDI-IMS

TMAs were imaged using a timsTOF Flex MALDI-QTOF mass spectrometer (Bruker Daltonics), images were collected at 200 laser shots per pixel in a mass range of 700-4000 m/z operating in a positive mode. Focus Pre TOF parameters were set as follows: transfer time 120.0 μs, pre-pulse storage 25.0 μs, and images were collected at 100um raster.

### MALDI-IMS Data Processing

Data sets were imported into SCiLs lab 2024a imaging software (Bruker) for data analysis. SciLs generated mass spectra that were normalized to the total ion count. Manual annotations for each m/z (mass-to-charge ratio) value were made based on an in-house database of confirmed N-glycan structures generated using GlycoWorkBench and GlycoMod for annotation[26,27]. In addition to MS-MS data previously done by our group[28]. The average intensity (peak maximum) of each m/z was extracted from SCiLs and used to calculate the relative intensity of each N-glycan m/z identified.

## Results

### Sialic acid isomers analysis in primary liver cancers

In this study, the AAXL method was applied to a paraffin-embedded tissue microarray (TMA) containing tissue from patients diagnosed with hepatocellular carcinoma (HCC), intrahepatic cholangiocarcinoma (iCCA), liver diseases like fatty liver, hepatitis, HPV (labeled here as others), and tissues from patients that had no sign of liver damage (labeled here as normal) and used for the identification and characterization of sialylated linkage specific N-glycan structures according to the sialic acid residue **(Figure 1a, and supplementary figure 1a)**. The difference in sialic acid linkage (α2,3, α2,6, and the combination α2,3 and α2,6) is illustrated by mass shifts seen in the image data **(Figure 1b)**. Since these linkages are considered isomers, data analysis of the mass will not resolve specific linkage, treatment with AXXL induces a mass shift in each type of linkage. α2,3-sialic acid linkage is identified by a +37.0316 Da mass shift and α2,6-sialic acid linkage is identified by a +27.0473 Da mass shift. Spectral mass shifts provide a clear identification between the two sialic acid isomers.

**Figure 1.**
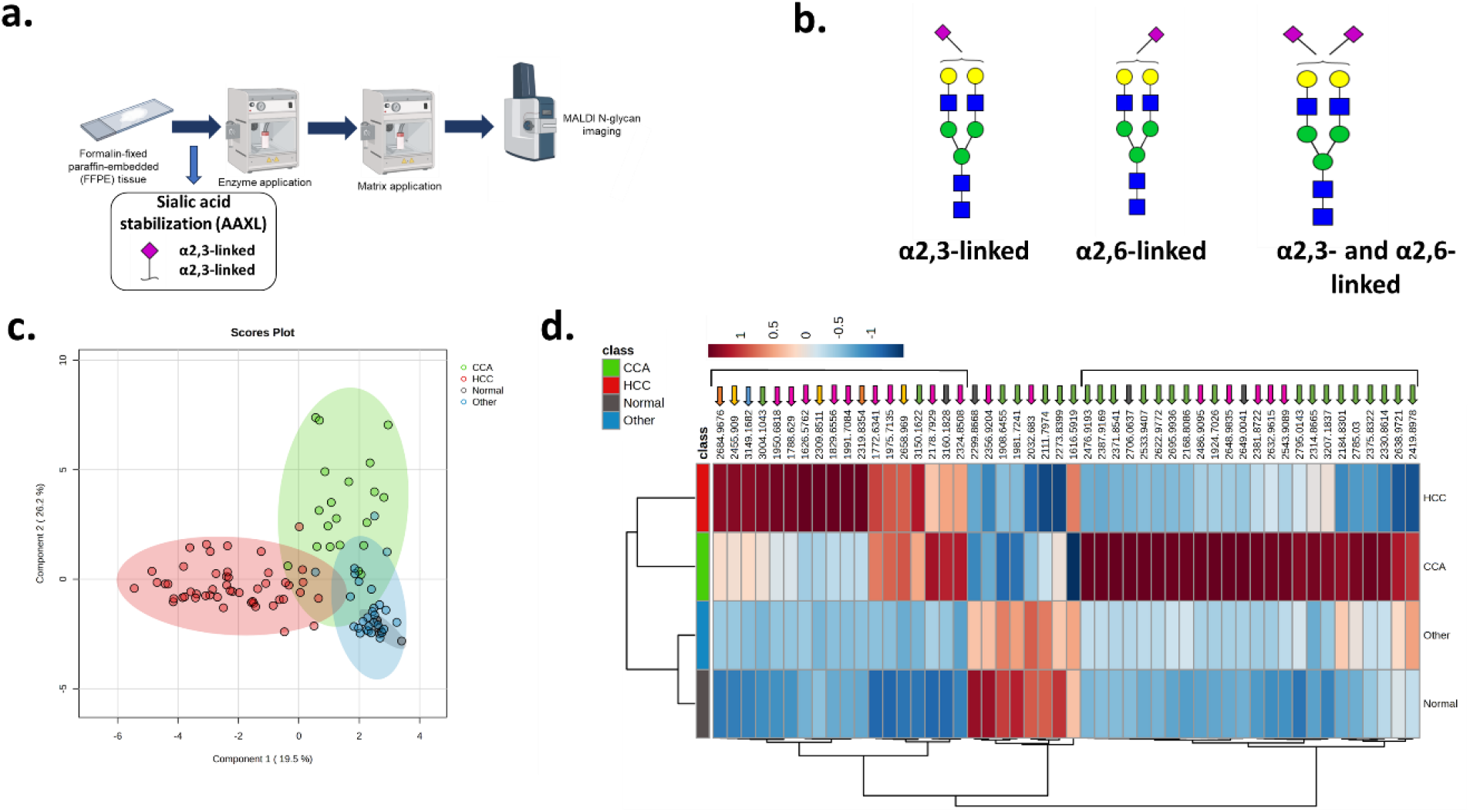
**a**. MALDI-IMS and sialic acid stabilization workflow outline. **b**. Sialylated N-glycan structure with labeled sialic acid linkages. Following the general nomenclature and illustration of N-glycans, sialic acid residues are represented by a purple diamond shape known to be attached to a galactose residue decorating the N-glycan structure. **c**. Clustering analysis of sialylated N-glycans based on respective groups. **d**. Heatmap analysis of top 50 N-glycan structure clustering based on respective groups. Colored arrows are representative of different linkages, pink: α 2-3, green: α 2-6, orange:2 α 2-3, yellow: α 2-3, α 2-6, gray:2 α 2-6, and blue: any other sialylated combination.

Global analysis of all sialylated N-glycans identified in the TMA revealed that sialylated N-glycans clustered different between iCCA and HCC groups **Figure 1c**. Similarly, heatmap analysis in **figure 1d** of the top 50 N-glycans demonstrates this significant relationship of specific N-glycan structures with respective groups. A clear trend was observed in the iCCA group where most of the N-glycans highly significant were those with an α2,6 sialic acid linkage while for the HCC group, there is a noticeable trend in N-glycans with an α2,3 sialic acid linkage. Normal and other tissues did not reveal specific trends but still had mixed sialic acid linkage patterns.

### Sialic acid linkages differ according to the type of liver cancer

Next, we analyzed our complete list of N-glycans identified from the TMA based on the type of sialic acid linkage (α2,3, α2,6, α2,3 & α2,6, 2 α2,3, and 2 α2,6). An α2,3 sialic acid linkage did not demonstrate a significant alteration between the types of cancer, only comparisons between each type of cancer relative to normal tissues had significant alterations **(Figure 2a)**. iCCA tissues demonstrated a significant α2,6 sialic acid linkage alteration relative to HCC tissues. α2,6 sialic acid linkages for HCC tissues were also decreased relative to normal tissues (**Figure 2b)**. However, for HCC the sialic acid linkage combination (α2,3 & α2,6) was significantly altered when compared to iCCA and normal tissues **(Figure 2c)**. Analysis of 2 α2,3, and 2 α2,6 linkages is presented in **supplementary figures 1c and 1d**, these did not demonstrate any significant trends of alterations between cancers. Here we demonstrate a clear difference in the type of sialic acid linkage with the type of liver cancer, where iCCA has more α2,6 linkages while HCC has more α2,3 & α2,6 linkages. Overall, this analysis establishes the value of identifying the specific sialic acid linkages according to the type of liver cancer for future studies.

**Figure 2.**
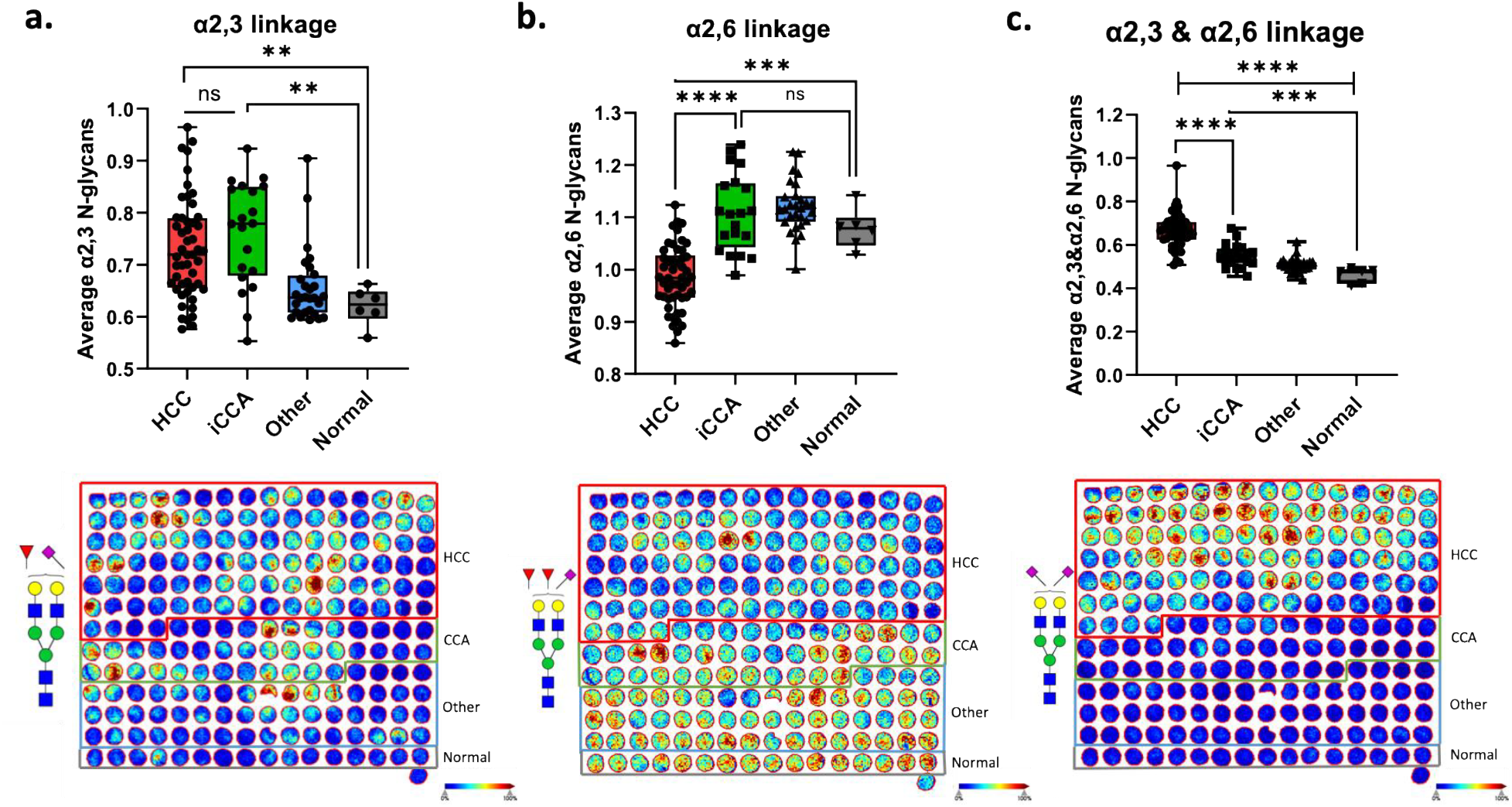
**a**. Box plot of the average of all N-glycans with an α2,3 sialic acid linkage (top), representative TMA images from α2,3 sialylated N-glycan and α2,3 sialylated, fucosylated N glycan (2137.766 m/z) (bottom). **b**. Box plot of the average of all N-glycans with an α2,6 sialic acid linkage (top), representative TMA images from α2,6 sialylated N-glycan and α2,6 sialylated, fucosylated N glycan (2273.839 m/z) (bottom). **c**. Box plot of the average of all N-glycans with an α2,3 & α2,6 sialic acid linkage (top), representative TMA images from α2,3/α2,6 sialylated N-glycan and α2,3/α2,6 sialylated, fucosylated N glycan (2309.851 m/z) (bottom).

### Fucosylated N-glycans with α2,6 sialic acid linkages are associated to iCCA

Previous studies by our group had identified fucosylated N-glycans to be highly altered in the tissue and serum of iCCA patients, we proposed the use of an N-glycan combination (including fucosylated N-glycans) to be a promising biomarker for detection of iCCA[29]. Thus, we were interested in identifying the same type of fucosylated N-glycans in this case with sialic acid decorations and determining if this specificity to iCCA maintains even with the attachments of sialic acid residues. **Figure 3a-c** shows representative MALDI-IMS images of non-sialylated N-glycan structures (similar as those previously reported): a glycan composed of Hex5dHex2HexNAc5 + 1Na (2158.780m/z), proposed to be bisected double fucosylated, a glycan composed of Hex5dHex1HexNAc5 + 1Na (2012.710 m/z, proposed to be a bisected single fucosylated, and a glycan composed of Hex4dHex2HexNAc5 + 1Na proposed bisected double fucosylated with one terminal galactose residue (1996.720 m/z).

**Figure 3.**
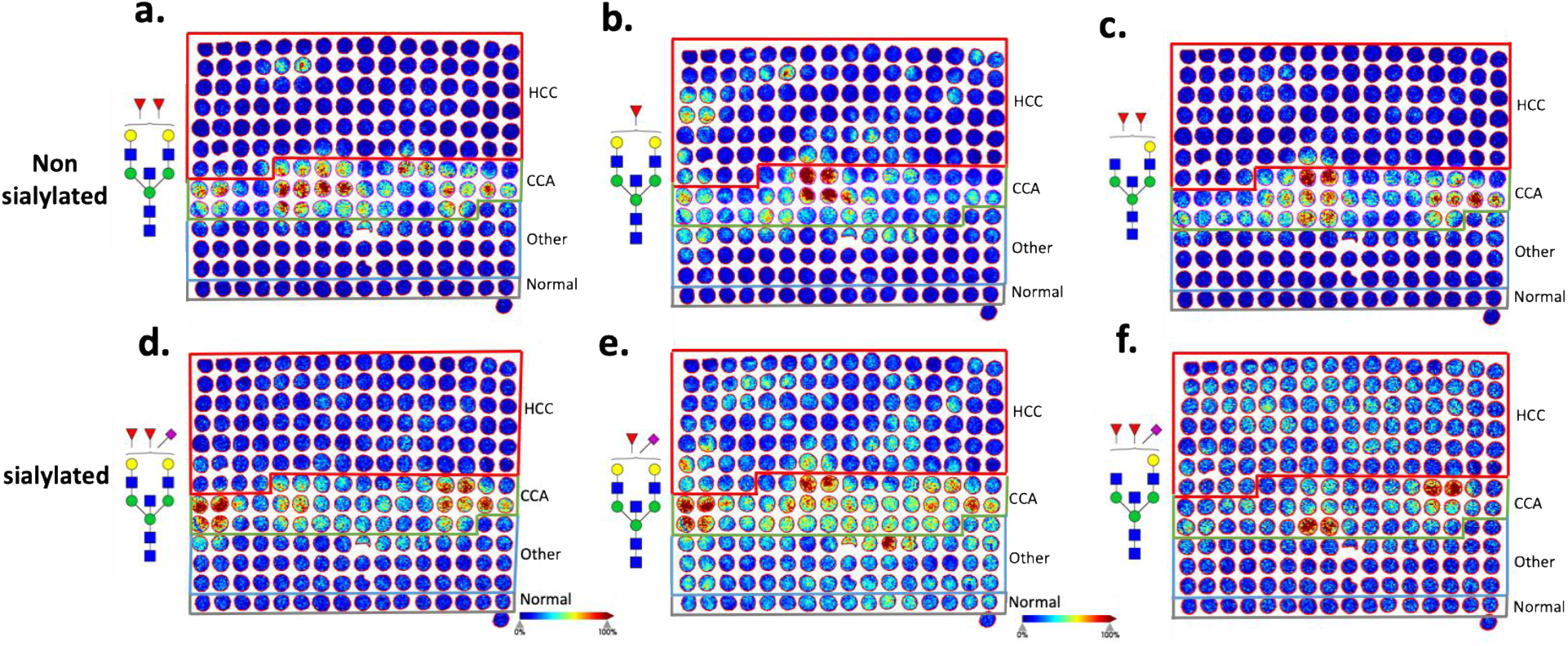
**a**. Representative TMA images from bisected fucosylated N-glycan. **b**. Representative TMA image from α2,6 sialylated, bisected fucosylated N-glycan. **c**. Representative TMA images from bisected fucosylated N-glycan. **d**. Representative TMA image from α2,6 sialylated, bisected fucosylated N-glycan. **e**. Representative TMA images from bisected fucosylated N-glycan. **f**. Representative TMA image from α2,6 sialylated, bisected fucosylated N-glycan.

**Figures 3d-f** show representative MALDI-IMS images of glycans with α2,6 sialic acid linkages. Figure 3d has the composition of Hex5dHex2HexNAc5 NeuAc1 + 1Na, proposed to be a singly sialylated, bisected an di-fucosylated N-glycan. Similarly, Figure 3e highlights that a glycan composed of Hex5dHex1HexNAc5 NeuAc1 + 1Na, proposed to be a singly sialylated, bisected fucosylated N-glycan. Similarly, Figure3f shows a glycan composed of Hex4dHex1HexNAc5 NeuAc1 + 1Na, proposed to be a singly sialylated, bisected di-fucosylated N-glycan with a single galactose residue. These α2,6 sialic acid linkage bisected fucosylated N-glycans not only maintained the same specificity but also had a higher intensity for some of the iCCA tissue cores relative to the non-sialylated structure.

### N-glycans differ according to the type of CCA

To further explore the presence of sialylated N-glycans in the context of CCA, we analyzed a second TMA for the N-glycan alterations (non-sialylated and sialylated) in different types of CCA (intrahepatic and extrahepatic). Extrahepatic CCA (eCCA) cores were divided according to the collection site of the tissue, that is bile duct (BD) and common bile duct (CBD) **(Supplementary figure 1b)**. As presented before in this study and our previous work (7), bisected double fucosylated N-glycan (2158.767 m/z) was significantly altered in iCCA patients and MALDI-IMS representative images for this cancer were highly specific. As **Figure 4a** demonstrates, a glycan composed of Hex5dHex2HexNAc5 + 1Na (2158.780m/z), proposed to be bisected double fucosylated glycan, is significantly altered in all types of CCA relative to normal tissues, no significant differences were observed within types of CCA. However, a glycan composed of Hex5dHex2HexNAc5 NeuAc1 + 2Na (2325.793/z), proposed to be a bisected, fucosylated and sialylated (α2,6 sialylated) N-glycan not only demonstrated a significant alteration in CCA types relative to normal but was also significantly altered in iCCA relative to the subtypes of eCCA, suggesting the importance of a sialic acid modification when differentiating between the types of CCA **(Figure 4b**). Other types of N-glycan structures like a high mannose glycan were highly altered in normal tissues relative to CCA types **(Figure 4c)**.

**Figure 4.**
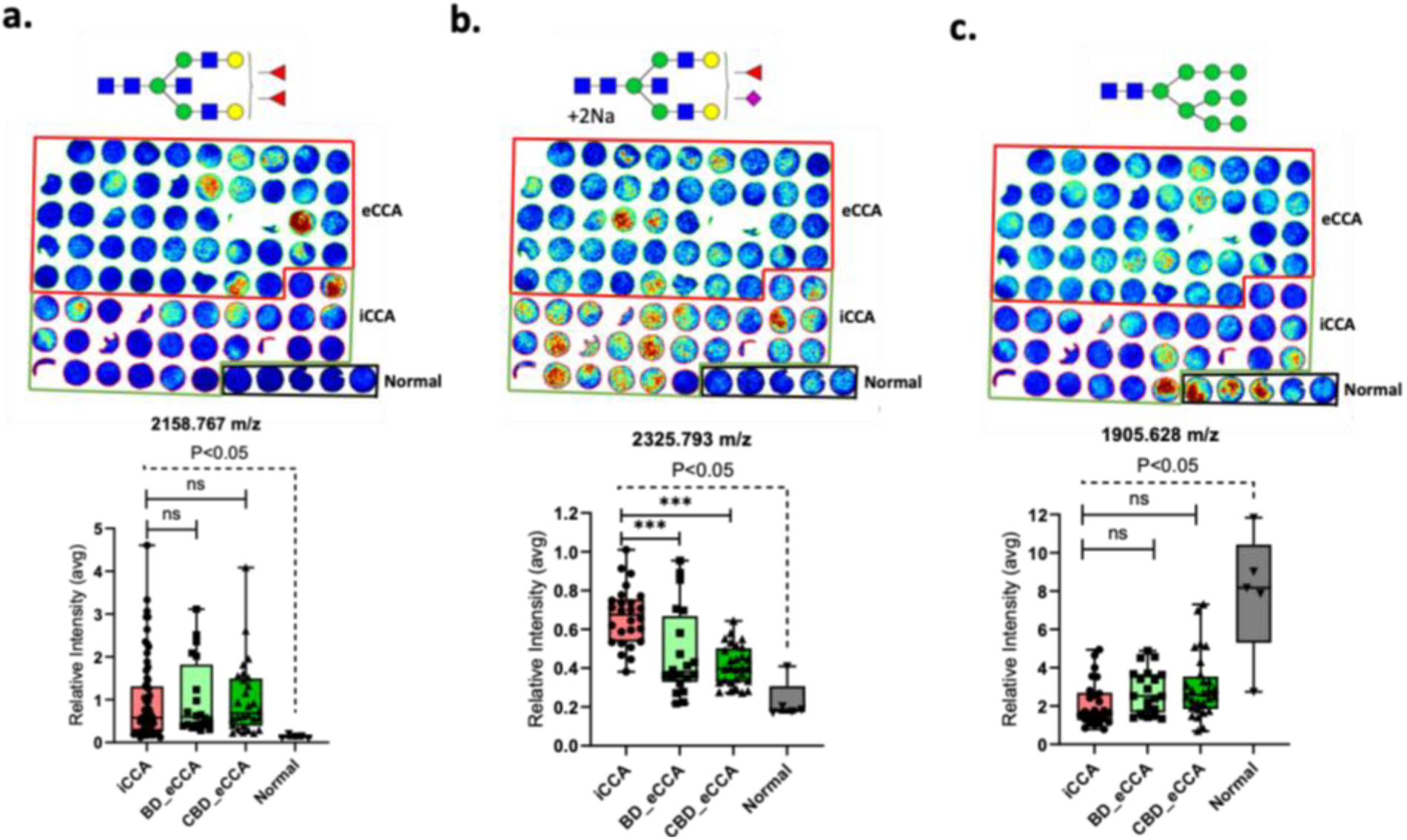
**a-c**. Representative TMA images from bisected fucosylated N-glycan (top). Relative Intensity quantification from respective N-glycan (bottom). BD: bile duct collection site, CBD: common bile duct collection site.

## Discussion

N-glycan sialyation on glycoproteins is a modification that is known to occur in cancer and has many biological roles. Since sialic acids are terminal modifications, these are known to interact with receptors on other cell surfaces, aid in cell signaling, and interact with siglec (sialic acid-binding Ig-type lectin) receptors for a role in immune modulation. α2,3 or α2,6 isomers are some of the most common modifications, both have been involved with different types of cancers.

Alterations in sialic acids, especially α2,6 linked sialic acid have been previously associated with several other cancers [16,18,19,30-32]. Indeed, there is some evidence that serum levels of α2,6 sialic acids may be altered in CCA [33]. However, up until now, no analysis has been performed directly in iCCA tissue. Indeed, previous methods to examine sialic acid through tissue N-glycan imaging were hampered by the method, which can be associated with sialic acid loss. Our recently described method stabilizes these sialic acids and allows for the determinations of linkage (α2,3 versus α2,6) [22].

Additionally, it is noted that CA19-9, the current “standard” biomarker for CCA, is a sialylated glycan with an α2,3-linked sialic acid. Previous reports by our group, identified bisected fucosylated N-glycans to be one of the major alterations to be highly specific in iCCA tissues and serum relative to any other type of liver disease, including HCC[29]. However, sialic acid residues were not explored in that study. Here, we address this gap and examine this modification and further identify specific linkages. This study elucidates the value of bisected fucosylated N-glycans in CCA and the bisected fucosylated sialylated, specifically for α2,6-linkage for CCA and CCA types. Future studies will extend this finding to a bigger tissue set cohort and determine if these N-glycan modifications can also be observed in serum and used for biomarker discovery.

In our initial work, we identified N-glycan alterations in iCCA tissue, and these were also found in the serum of patients with iCCA [29]. Here we have extended that work to other forms of CCA, specifically, both pCCA and dCCA. The major glycan alteration found is proposed to be a bisected doubly-fucosylated bi-antennary glycan [29]. Similarly, a paper by an independent group has confirmed this finding specifically in iCCA and shown that this glycan modification sustains the NOTCH and EGFR/NF-κB signaling pathways and has a prognostic value in iCCA[34]. This independent finding supports our result and highlights the value of leveraging this information as a biomarker of CCA.

In conclusion, we have used MALDI-IMS to identify specific alterations in N-linked glycosylation that are associated with defined subtypes of CCA. As we have already shown that the N-glycans found in tissue are also reflected in the circulation, it is hoped that future work will identify the glycoproteins that contain these glycan changes and leverage those to help in the early diagnosis of this deadly cancer.

