## Supplemental figure 1 for "Identification of sialic acid linkage profiles in primary liver cancers"

**Corresponding Author:**

Anand S. Mehta

Medical University of South Carolina

**Supplementary Figure 1**

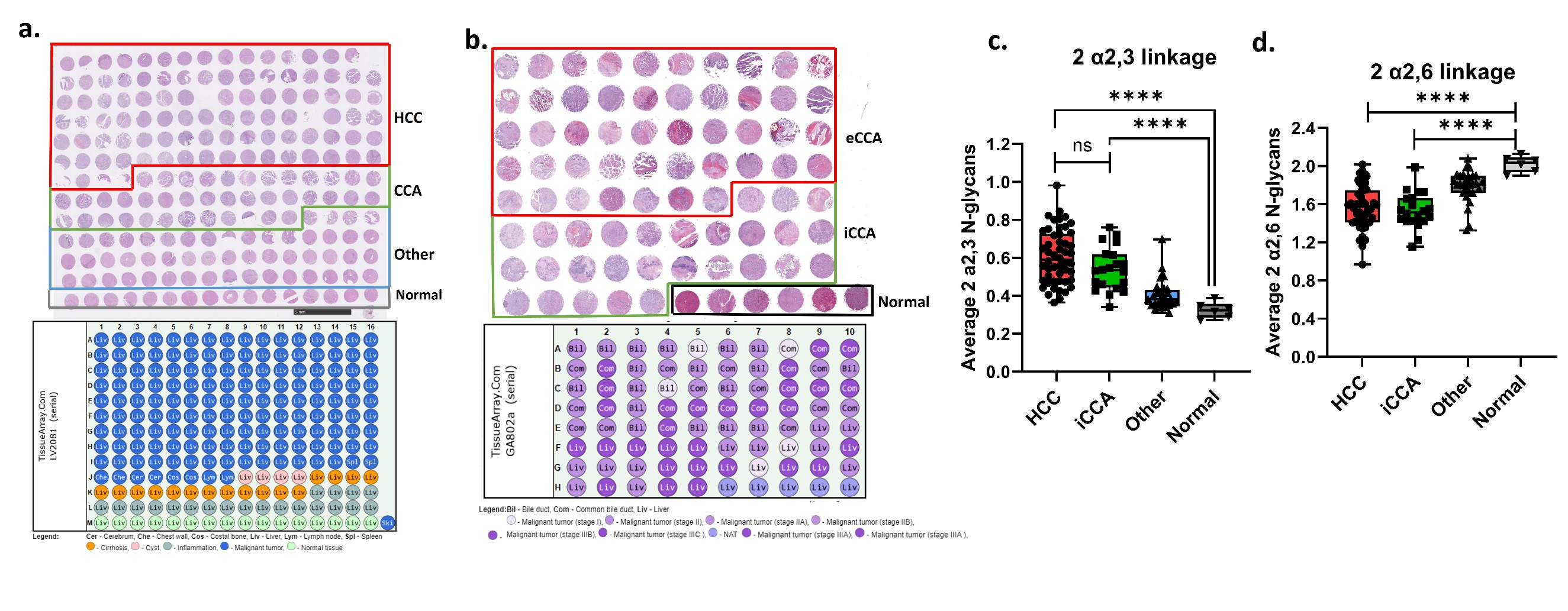
**Supplemental Figure 1.** Hematoxylin & Eosin (H&E) stainings with respective outline of specimens included for tissue microarrays (TMAs). (a) TMA 1; #LV2081 and (b) TMA 2; #GA802a. c. Box plot of the average of all N-glycans with 2 α2,3 sialic acid linkages (left) and 2 α2,6 sialylated N glycan.
